# Benchmarking large language models for ACMG/AMP variant interpretation and variant calling

**DOI:** 10.64898/2026.06.30.735646

**Authors:** Manuel Corpas

**Affiliations:** Faculty of Life Sciences, University of Westminster, London, UK

**Keywords:** agentic AI, large language models, variant interpretation, variant calling, ACMG/AMP, ClinVar, Genome in a Bottle, benchmarking, trustworthy AI, clinical genomics

## Abstract

Agentic large language models are increasingly used across the genomic workflow, from variant calling to clinical interpretation, yet they are evaluated by accuracy alone, a single figure that cannot say whether a system is safe or where in the workflow a failure originates. We present ClawBench, a framework that attributes each outcome to the architectural layer that produced it across both halves of the canonical pipeline. Two design choices remove the confounds that make agentic genomics hard to evaluate: a temporally blinded truth set, in which every scored ClinVar label first became available only after the training cutoff of every model tested, and a fail-closed evidence contract that blocks evidence circular with the truth label. We score validity, safety, provenance and reproducibility, not accuracy alone, under a constraint gradient that relocates correctness from a model’s prior into executed, validated code.

We show three things. First, dangerous misclassification is rare and model-invariant, a controlled precondition of the executed architecture rather than a frontier, while fabricated evidence is measurable and is neutralised by execution. Second, different variant classes are rate-limited by different layers: loss-of-function variants by the deterministic combiner threshold, and rare missense by evidence formation, where evidence acquisition is asymmetric and capped and strength assignment is a recoverable layer that naive strength-licensing prompts confound. Third, for variant calling the arms separate not on whether a model can plan a pipeline, which all do, but on trust properties, pinning, provenance, auditability and reproducibility, which climb monotonically toward validated execution; and a local open-weight model reproduces the safety result yet meets the structured-output and provenance contract far less often than frontier models, a conformance gap rather than a capability or safety gap. An end-to-end join attributes failures across the whole workflow, separating a missed call from a propagated genotype error from a correctly called but misinterpreted variant.

ClawBench shows that apparently identical outcomes arise from distinct, independently measurable failure modes, and that trustworthiness in agentic genomics is a property of the pipeline architecture rather than of the model, providing a portable, contamination-resistant unit of attribution for the field.

## Introduction

Large language models are moving from answering medical questions to executing clinical tasks, and clinical variant interpretation is among the most consequential [1]. Classifying a germline variant under the American College of Medical Genetics and Genomics and Association for Molecular Pathology (ACMG/AMP) framework [2] requires assembling many lines of evidence, population frequency, computational prediction, segregation, functional data, and combining them into one of five tiers from pathogenic to benign. Agentic systems that retrieve evidence, assign ACMG criteria and return a classification are now being built and benchmarked for exactly this task [3]. The clinical stakes are high: a misclassified variant can withhold a needed intervention or trigger an unnecessary one.

Yet the way these systems are evaluated cannot tell us whether to trust them, for two reasons. First, evaluation is almost always reduced to accuracy, the fraction of variants whose final tier matches a reference. A single accuracy figure cannot distinguish a system that is safe but uncertain from one that is confidently wrong, and it cannot say why a given variant was misclassified. Recent benchmarks already show the symptoms that an accuracy score hides: models classify variants with strong evidence well but become inconsistent when evidence is weak, and they tend to overclassify, assigning evidence at higher strength than warranted [3]. In the clinic such confident error is the dangerous failure mode, and language models are known to produce plausible, fluent fabrications that are difficult to detect [4]. Second, two confounds make naive benchmarking on public knowledge bases unsound. A model may have memorised a variant’s published classification during pretraining, so any test drawn from a resource such as ClinVar [5] risks measuring recall rather than reasoning [6]; and evidence sourced from the same database that supplies the truth label is circular, inflating apparent competence. Variant interpretation is genuinely hard even for expert laboratories: applying the ACMG/AMP guidelines to the same 99 variants, nine laboratories reached only 34 per cent concordance before consensus review, rising to 71 per cent afterwards [7]. A benchmark that conflates this intrinsic difficulty with model behaviour, and that cannot rule out contamination or circularity, cannot support a trust decision.

The deeper limitation is conceptual. An agentic interpretation pipeline is a stack of distinct steps, retrieving evidence, deciding which criteria apply, choosing how strong each is, and combining them, and a failure can occur at any one of them. A scalar score collapses this structure into a single number, so a variant returned as uncertain significance carries no indication of whether the uncertainty came from missing evidence, an unstable choice of criteria, a miscalibrated strength, or a combining threshold. What is missing is not a better score but a different unit of analysis: an attribution of each variant’s outcome to the architectural layer that produced it.

We present ClawBench, a framework that provides this attribution. It rests on two methodological foundations that remove the confounds above: a temporally blinded truth set, in which every scored label first became available only after the training cutoff of every model under test, and a fail-closed evidence contract that blocks evidence whose provenance is the truth label. Within this design, each variant is interpreted under a constraint gradient that progressively moves correctness from the stochastic model into executed, validated code, and the resulting structured submissions are mapped to an ordered stack of architectural layers grouped into three blocks: safety control; evidence formation, comprising acquisition, sufficiency, assignment and strength calibration; and decision formation. The framework assigns every variant a layer-attribution label rather than a score. Using it, we show that safety is a controlled, model-invariant precondition, that fabricated evidence is measurable and is neutralised by execution, and, crucially, that different variant classes are rate-limited by different layers: loss-of-function variants by the combining threshold, and rare missense by evidence formation. Two targeted experiments then dissect the missense frontier. An oracle acquisition arm shows that supplying complete, real, non-circular evidence is a genuine but asymmetric and capped intervention, and a deterministic strength-calibration re-score, confirmed on a second model, shows that the residual frontier is strength assignment, a partially recoverable layer whose naive instruction-based licensing is prompt-fragile. This work builds on and complements a companion pharmacogenomics benchmark [8], which established that interpretation trustworthiness is a property of pipeline architecture rather than of the model; here we decompose that architecture into independently measurable layers and show that apparently identical classification outcomes arise from fundamentally different, and separately diagnosable, failure modes.

## Results

Across the pilot, exact five-tier agreement with the held-out truth was similar across the constraint-gradient conditions and the three models (roughly 48 to 55 per cent; Table 1). A single accuracy figure therefore hides, rather than explains, the behaviour of these systems. The results below instead attribute each variant’s outcome to a specific architectural layer, and show that the layer carrying the uncertainty differs by variant class.

**Table 1:**
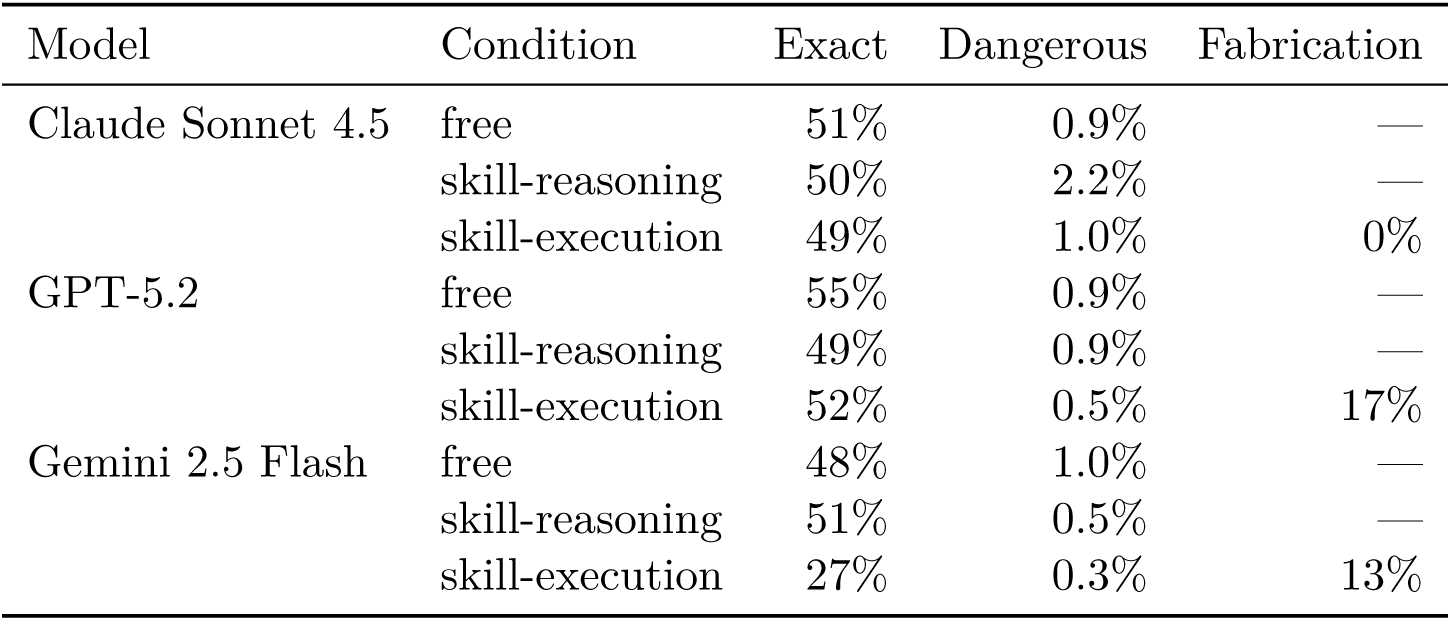
Tier-A pilot, by model and constraint-gradient condition (231 variants, five replicates). Exact: five-tier agreement with truth. Dangerous: Pathogenic-to-Benign miscall rate. Fabrication: fraction of skill-execution submissions with at least one ClinVar-derived code stripped by the contract.

### A contamination-resistant, tiered held-out benchmark

Temporal blinding admitted 6,929 of 166,437 two-star GRCh38 candidates (206 reclassified); the fail-closed first-availability filter excluded 159,300 pre-cutoff records and 208 with undatable history. Every admitted label first became available on or after 2025-11-29, at least 91 days after the binding model cutoff (GPT-5.2, 2025-08-31). The set is immutable and content-hashed, reproduces bit-for-bit, and spans the automatability spectrum (Tier A 894, Tier B 2,906, Tier C 3,129), so any measured difference localises to a layer rather than to set composition. These guarantees apply to the benchmark *artefacts*, the frozen truth set and the deterministic combiners; model behaviour is stochastic and is reported over five replicates, with agreement on the modal class ranging from 37 to 97 per cent across the model-by-condition cells (lowest for the most uncertain free-prompted cell, highest under skill-reasoning).

### Safety is model-invariant and a controlled precondition

The lethal failure mode, calling a benign variant pathogenic or the reverse, was rare in every condition and every model. In the Tier-A pilot the dangerous-miscall rate ranged from 0.3 to 2.2 per cent across the nine model-by-condition cells (Table 1). Across the full skill-execution attribution corpus, the safety-clean flag held for 99 to 100 per cent of variants in every consequence group (loss-of-function, missense and other). Dangerous misclassification is therefore rare (0.3 to 2.2 per cent) and model-invariant rather than a frontier; we describe it as a controlled precondition rather than “solved”, since a 2.2 per cent pathogenic-benign confusion rate is not clinically negligible at scale, and it is a property of the executed architecture, not of the individual model.

### Fabricated evidence is measurable and neutralised by execution

Under skill-execution the contract strips evidence whose provenance is the truth label before scoring. The fraction of submissions containing at least one such fabricated, ClinVar-derived code was strongly model-dependent: 17 per cent for GPT-5.2, 13 per cent for Gemini 2.5 Flash and 0 per cent for Claude Sonnet 4.5. The execution layer converts this fabrication into safe abstention rather than a dangerous call: when Gemini’s fabricated codes were stripped it fell back to uncertain significance, lowering its exact-match accuracy to 27 per cent while keeping its dangerous-miscall rate at 0.3 per cent. Confident circular reasoning is thus rendered measurable and defused, not silently scored as correct.

### Pathogenic versus Likely-Pathogenic disagreement is a combiner-threshold effect

Apparent disagreement between the Pathogenic and Likely-Pathogenic tiers is predominantly an artefact of the deterministic combiner threshold, not of evidence. Re-scoring identical model-assigned codes through the Tavtigian point engine [9] instead of the Richards rule [2] flipped the modal class for 74 per cent of loss-of-function variants, almost entirely at the Likely-Pathogenic to Pathogenic boundary (349 of the combiner-sensitive transitions), reflecting the PVS1-plus-PM2 configuration that the rule caps at Likely Pathogenic but the points total scores as Pathogenic. We treat PVS1 atomically at its default strength here; its own loss-of-function strength decision tree [10] could refine attribution within this class further and is a natural extension for a loss-of-function-dominated analysis. This divergence between the Richards rule and the Tavtigian points system for the PVS1-plus-PM2 configuration is a documented property of the two scoring frameworks, not a behaviour of the agents: the models assigned the evidence correctly, and the disagreement is a property of the combiner choice. Combiner sensitivity is therefore a decision-formation phenomenon concentrated in loss-of-function variants.

### Missense uncertainty is dominated by evidence sufficiency

The same comparison shows a different bottleneck for rare missense. Combiner sensitivity affected only 6 per cent of missense variants, and where present it sat at the uncertain-significance to Likely-Benign boundary, not the pathogenic boundary. Instead, 55 per cent of missense variants were evidence-insufficient: the truth was definitive, yet even the point engine returned uncertain significance because molecular consequence and allele frequency alone do not constitute enough evidence to classify. The dominant layer for missense is therefore evidence formation (sufficiency and acquisition), whereas for loss-of-function it is the combiner threshold. Different variant classes are rate-limited by different layers.

### Acquisition is a real but asymmetric and capped layer

We then asked whether giving an agent complete, real, non-circular evidence resolves missense uncertainty. On 24 definitive rare-missense variants, supplying oracle evidence (Ensembl VEP annotations [11] including calibrated REVEL [12] and AlphaMissense [13]) moved 7 of 24 variants out of the evidence-insufficient category (29 per cent), versus 24 of 24 evidence-insufficient under thin evidence. The effect was entirely one-directional (Table 2): all seven resolutions were on the benign side, each via a calibrated BP4 at supporting strength that reaches Likely Benign on its own, while none of the twelve pathogenic-side variants resolved. Three uncertain-significance negative controls correctly remained uncertain. Acquisition is thus a genuine layer, it moves variants that sufficiency alone could not, but it is asymmetric and capped: even with complete oracle evidence, the theoretical ceiling for these variants is 9 of 24, because the non-circular evidence available for a rare pathogenic missense (PM2 plus a calibrated PP3) reaches Likely Pathogenic only when PP3 attains strong strength.

**Table 2:** Acquisition arm (Claude Sonnet 4.5, 24 definitive rare-missense variants). Evidence-insufficient counts under thin versus oracle-enriched evidence, by truth class.

| Truth class | n | Thin | Enriched |
| --- | --- | --- | --- |
| Pathogenic | 6 | 6 | 6 |
| Likely Pathogenic | 6 | 6 | 6 |
| Likely Benign | 6 | 6 | 2 |
| Benign | 6 | 6 | 3 |
| Total | 24 | 24 | 17 |

### Strength assignment is a recoverable layer, and naive licensing is prompt-fragile

The gap between the realised 17 of 24 evidence-insufficient and the 9 of 24 ceiling is an assignment-strength effect: Claude Sonnet 4.5 applied PM2 at supporting (the SVI-recommended strength) in 101 of 101 instances, one point short of the Likely-Pathogenic threshold for the strong-PP3 pathogenic variants. To isolate this strength variable within a single convention, we re-scored the model’s own enriched-arm code sets deterministically, holding every non-PM2 code at its submitted calibrated strength and moving only PM2 from supporting to its moderate baseline. Under this consistent comparison, raising PM2 to moderate recovered two pathogenic variants and regressed no benign ones, in both models (evidence-insufficiency 17 to 15 for Claude Sonnet 4.5 and 18 to 16 for GPT-5.2; Table 3). The absence of a benign cost is mechanistic rather than empirical: the benign-side variants do not carry PM2, so altering its strength is inert there, and moderate-PM2 was therefore not tested against benign signal and found harmless. For the same reason the two-model agreement is a confirmation of an arithmetic property of each model’s submitted code sets, not an independent empirical replication. Strength assignment is therefore a real layer that is recoverable by a deterministic post-hoc re-score, an architectural intervention rather than the model behaving better when prompted: when the identified code set is held fixed, the residual closes in the pathogenic direction at no benign cost.

**Table 3:** Strength assignment, isolated by a deterministic consistent-convention re-score (PM2 supporting versus moderate, all other codes held at their submitted calibrated strength) on 24 definitive rare-missense variants. Moving PM2 to moderate recovers pathogenic variants at no benign cost in both models; the benign regression seen under a re-prompting protocol was an instruction-induced artefact, not a property of the arithmetic.

| Model | Recovered (pathogenic) | Regressed (benign) | Insufficient (PM2 sup. → mod.) |
| --- | --- | --- | --- |
| Claude Sonnet 4.5 | 2 | 0 | 17 → 15 |
| GPT-5.2 | 2 | 0 | 18 → 16 |

An earlier protocol that instead re-prompted the model to license PM2 at moderate appeared to show a direction-coupled trade-off, with benign resolutions regressing and net insufficiency rising to 22 of 24. That trade-off was an artefact of the instruction, not of the arithmetic. Licensing PM2 caused the model to apply it to benign variants where it had previously and correctly omitted it, raising PM2 use on benign-side variants from 27 of 60 to 59 of 60 replicates; the over-applied pathogenic-direction code, not its strength, produced the regression. The deterministic re-score, which alters only the strength of the codes the model actually identified, removes this confound. The finding is therefore twofold: strength assignment is recoverable under a consistent convention, and naive strength-licensing prompts are fragile, since a single instruction leaks into code selection, a confound an attribution framework must itself control for.

### Per-variant attribution localises uncertainty to architectural layers

Taken together, the framework assigns every variant an attribution label rather than a score, and those labels are not interchangeable: an uncertain-significance outcome is variously safety-clean, evidence-insufficient, assignment-unstable, calibration-sensitive or combiner-sensitive, and the benchmark reports which. The dominant layer is a property of the variant class (Table 4): loss-of-function variants are rate-limited by the combiner threshold; rare missense by evidence sufficiency, then assignment, then strength calibration; and safety is a controlled precondition across all classes. Apparently identical classification outcomes therefore arise from distinct architectural failure modes, which this framework attributes and measures independently. Because the attribution is computed from a model’s structured submissions against a fixed contract, any laboratory can run it on its own agentic pipeline to learn which layer limits its variants, independent of the underlying model.

**Table 4:** Dominant rate-limiting layer by variant class (per-variant attribution across the skill-execution corpus).

| Variant class | Dominant layer |
| --- | --- |
| Loss-of-function | Combiner threshold (74% combiner-sensitive) |
| Rare missense | Evidence sufficiency (55%), then assignment and strength calibration |
| All classes | Safety controlled (99–100% safety-clean) |

### Execution: trust is a property of the architecture, not the model

The interpretation results above concern what a model concludes. We next ask whether an agent can be trusted to *execute* the upstream genomic workflow, calling germline short variants from sequencing reads, under the same constraint gradient. The gradient is the same in structure (free, skill-reasoning, skill-execution and a reference control) but realised on a different axis: in interpretation each arm ends in a deterministic ACMG combiner, whereas in calling each ends in a containerised pipeline, so the layer being relocated is the calling toolchain rather than the evidence combiner. Four arms call the same GIAB benchmark sample: a free agent given only the task, a skill-reasoning agent given the wrapper specification, a skill-execution agent that invokes the validated pinned wrapper, and a best-practice reference configuration. Each run receives a set of pre-registered failure labels rather than a single verdict, by a deterministic classifier (Methods).

Two findings separate the arms, and neither is accuracy. First, plan validity does not discriminate: every arm, free and skill-guided alike, emitted a workflow that ordered the mandatory stages correctly, the free agent by hand-authoring alignment then calling and the skill arms by invoking a complete pipeline. Planning, like safety in interpretation, does not discriminate the arms for current frontier models on this task. Second, the trust properties do discriminate, and monotonically (Table 6). Moving from free prompting to skill-reasoning to validated execution, the fraction of runs that are container-pinned rises from none to full, provenance emission rises from none to complete, and auditability and reproducibility follow. The free agent authored plausible workflows that were unpinned, unprovenanced and not guaranteed reproducible; the validated arm was clean on every dimension by construction.

**Table 5:**
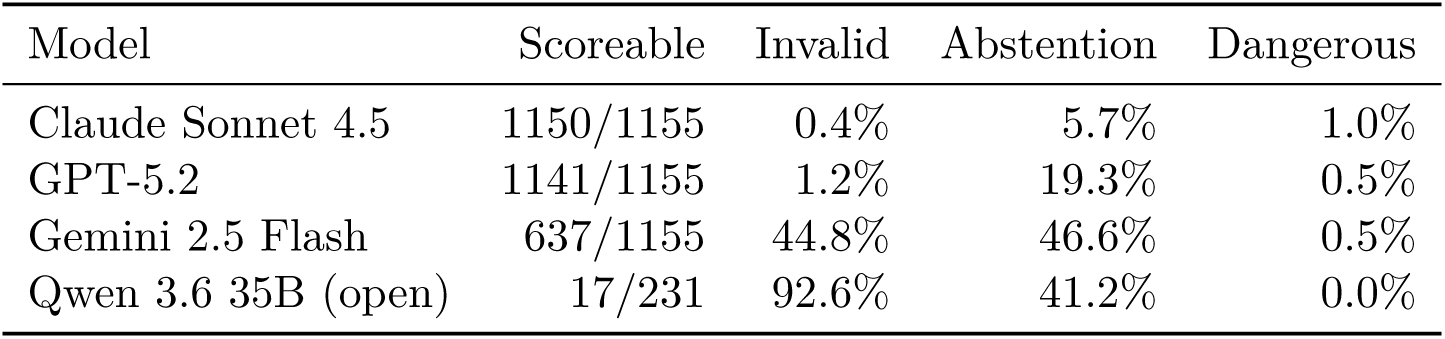
Open-weight versus frontier under skill-execution (same 231 temporally blinded Tier-A variants; one replicate for the open arm, five for the frontier arms). The open model emits parseable JSON and is safe, but the fail-closed contract rejects most of its submissions on structured-output and provenance grounds (absent confidence field, empty source identifier), not on classification. Invalid rate, not abstention, is the discriminating axis.

| Model | Scoreable | Invalid | Abstention | Dangerous |
| --- | --- | --- | --- | --- |
| Claude Sonnet 4.5 | 1150/1155 | 0.4% | 5.7% | 1.0% |
| GPT-5.2 | 1141/1155 | 1.2% | 19.3% | 0.5% |
| Gemini 2.5 Flash | 637/1155 | 44.8% | 46.6% | 0.5% |
| Qwen 3.6 35B (open) | 17/231 | 92.6% | 41.2% | 0.0% |

**Table 6:** Execution constraint gradient (GIAB HG002, chr20). Trust properties, derived from the emitted workflow text, climb across the gradient while plan validity is constant. Calling F1 is reported as qualification of the validated control arm, not as a result.

| Arm | Pinned | Provenance | Auditable | Plan valid | F1 |
| --- | --- | --- | --- | --- | --- |
| Free agent | no | no | no | yes | n/a |
| Skill-reasoning | partial | yes | no | yes | n/a |
| Skill-execution (validated) | yes | yes | yes | yes | 0.99, reproducible |

The validated arm’s accuracy is reported as qualification, not as a result: on GIAB chr20 it reached an F1 of 0.99 with a genotype-identical re-run, the expected behaviour of a pinned, validated pipeline. The contribution is the contrast in trust properties, which a single accuracy figure cannot express: an unpinned, unprovenanced workflow that happens to produce a plausible variant call is not a trustworthy clinical artefact, and the framework records exactly why.

### Equity is a property of the pipeline and is reported honestly

Stratifying the validated arm by GIAB sample ancestry, the calling F1 showed no large difference across the three ancestries tested (European, Ashkenazi, East Asian; spread 0.04 per cent). This F1 stability is not ancestry invariance, and we do not claim it as such. The same data carry a precision and false-positive-rate gradient roughly eight times the F1 spread, with the East Asian sample carrying the highest false-positive burden (about 32 per cent more false positives per true positive than the Ashkenazi sample), and the European and East Asian precision confidence intervals do not overlap (Table 7). These intervals treat each called variant as independent, whereas the ancestry contrast rests on one to two genomes per group, so the effective sample for an ancestry-level statement is the individual, not the variant; the difference is better read as a between-sample observation confounded with coverage and library preparation than as a population-level precision estimate. We therefore claim only that calling F1 does not differ materially across these three GIAB samples, and we treat this as a scope limitation rather than an equity result: any property here is a property of the validated pipeline and the GIAB resource, not of the agentic layer, and GIAB does not include African, South Asian or Indigenous American genomes, so ancestry coverage is narrow.

**Table 7:** Calling performance by GIAB sample ancestry (chr20, counts pooled within ancestry; Wilson 95% intervals). F1 is near-flat, but precision and the false-positive rate are not, and the East Asian and European precision intervals do not overlap. The ancestry-invariance claim is withdrawn.

| Ancestry | *n* | Precision (95% CI) | Recall | F1 | FP per TP |
| --- | --- | --- | --- | --- | --- |
| European | 1 | 0.9896 (0.9889–0.9903) | 0.9920 | 0.9908 | 1.05% |
| Ashkenazi | 2 | 0.9893 (0.9888–0.9898) | 0.9917 | 0.9905 | 1.08% |
| East Asian | 1 | 0.9863 (0.9855–0.9871) | 0.9944 | 0.9903 | 1.39% |

### An open-weight model reproduces safety but not the provenance contract

A contamination-resistant benchmark should not depend on a paid endpoint, so we ran the full interpretation gradient on a local open-weight model (Qwen 3.6 35B, served on-device with frozen weights) over the same 231 temporally blinded Tier-A variants as the frontier models, at one replicate. The result separates safety from conformance. The open model reproduced the safety finding: it emitted schema-parseable JSON on every call (format-failure rate zero), its dangerous-miscall rate was at most 1.7 per cent across the three arms (zero among the execution-scored variants), and its free-prompting and skill-reasoning arms were fully scoreable (231 of 231), with substantively sensible ACMG reasoning (for a common benign variant it proposed BS1 and BP4 with correct rationale).

Under skill-execution, however, the fail-closed contract rejected 214 of its 231 submissions as invalid (92.6 per cent), against 0.4 per cent for Claude Sonnet 4.5, 1.2 per cent for GPT-5.2 and 44.8 per cent for Gemini 2.5 Flash (Table 5). The rejections were not classification errors, since the combiner derives a class from codes and strengths alone and these were largely correct; they were structured-output and provenance failures. Of the 214, 168 omitted the per-code confidence field the schema requires and 169 left the source identifier empty, with a few cases over-strengthening a benign code or citing an inadmissible source type. The empty source identifier is the auditability property the framework measures, so we report the gap as observed rather than populating it, which would inflate the very trust property under test. The open model therefore localises its deficit to the trust contract, structured provenance and auditability, not to safety and not to classification reasoning, and Gemini 2.5 Flash sits on the same gradient as a weak-frontier exemplar. The open-versus-frontier contrast is thus one of structured-output and provenance conformance, a property the framework was built to expose, and it is established on free, on-device weights that a paid-API-only benchmark could not. This was a single-replicate pass; a powered multi-replicate run, and whether stronger schema elicitation closes the conformance gap, are future work.

### An end-to-end join attributes failure across the whole workflow

Finally we connect the two halves. For variants that are both GIAB-confident truth calls and members of the held-out interpretation set, the framework joins each variant’s calling outcome to its interpretation attribution, yielding a single end-to-end label: a missed call (calling layer), a genotype error that propagates into interpretation (cross-layer), a correctly called variant rate-limited by a named interpretation layer, or a dangerous misclassification (the safety apex). On the GIAB samples this overlap is small and entirely benign (16 distinct chr20 variants, all correctly called, all benign), which is itself a finding: reference genomes from healthy individuals are not enriched for pathogenic variants, so the natural call-and-interpret overlap is benign-dominated, and pathogenic end-to-end cases require clinical cohorts or controlled spike-ins, which we leave to future work. The value demonstrated here is the unit of attribution itself: where a scalar end-to-end accuracy records every failure identically as “wrong”, the join assigns a distinct, actionable label (a missed call, a propagated genotype error, a named interpretation-layer limit, or a dangerous misclassification) that localises each failure to a specific workflow layer. On the benign-only GIAB overlap no variant exercises these failure labels; we therefore present the join as a validated mechanism with the label taxonomy it produces, and reserve a populated demonstration for a cohort with genuine call-and-interpret failures.

## Discussion

Accuracy is the wrong primary endpoint for agentic genomics. A single figure cannot say whether a system is safe, and it cannot say where in the workflow a failure arose. ClawBench replaces the figure with a unit of attribution: every outcome, whether a variant call or a clinical interpretation, is assigned to the architectural layer that produced it. Two results recur across both halves of the workflow and together define the contribution. Relative to the companion pharmacogenomics benchmark [8], which established that interpretation trustworthiness is a property of pipeline architecture rather than of the model, the advance here is the machinery that makes that claim diagnosable rather than asserted: provable temporal blinding with a fail-closed first-availability filter, per-layer attribution as a portable unit, the dissection of the rare-missense frontier into acquisition and strength-assignment layers, and the finding that naive strength-licensing prompts leak into code selection.

First, the property that matters is not capability but architecture. In interpretation, safety is a controlled, model-invariant precondition; in execution, plan validity holds for current frontier models. What separates trustworthy from untrustworthy systems is not whether the model can classify a variant or order a calling pipeline, which they largely can, but whether the surrounding architecture enforces non-circular evidence, pins its tools, records provenance and reproduces. Trustworthiness migrates from the stochastic model into the executed, validated code. This is why a constraint gradient, rather than a leaderboard, is the right instrument: it relocates correctness layer by layer and measures what each layer is worth.

Second, apparently identical outcomes have distinct, measurable causes. An uncertain-significance call may be safety-clean, evidence-insufficient, assignment-unstable, calibration-sensitive or combiner-sensitive, and the dominant layer is a property of the variant class: loss-of-function variants are rate-limited by the combiner threshold, rare missense by evidence formation, where acquisition is asymmetric and capped and strength assignment is recoverable by a deterministic re-score of a fixed code set, while naive prompt-based licensing of that strength is confounded by prompt-sensitive code selection. The end-to-end join extends the same logic across the pipeline, distinguishing a missed call from a propagated genotype error from a correctly called but misinterpreted variant. A laboratory can run the framework on its own agentic pipeline, with a frontier or a local open-weight model, to learn which layer limits its variants.

The framework is deliberately contamination-resistant and reproducible. Temporal blinding ensures no tested model could have seen a scored label during pretraining, the fail-closed contract blocks truth-circular evidence, and the open-weight arm makes the interpretation benchmark runnable on-device with frozen, hash-identifiable weights. These choices matter more for an evaluation standard than any single score, because a benchmark that cannot be reproduced, or whose labels leak into the next model, measures nothing durable.

### Limitations

This study is scoped to the measurement framework, and several boundaries are stated rather than hidden. Calling accuracy is reported as qualification of the validated control arm on a chromosome-20 development pilot; we do not claim a genome-wide calling benchmark, and the calling F1 is the expected behaviour of a validated pipeline rather than a finding. Reproducibility is demonstrated as same-host genotype-identical re-runs; cross-host and clean-rebuild reproducibility on multiple samples remain future work. The equity analysis withdraws any ancestry-invariance claim: GIAB spans only three ancestries and is benign-dominated, the East Asian sample carries a higher false-positive burden than F1 alone reveals, and broader population coverage requires cohorts beyond GIAB. The end-to-end pathogenic cases are exercised through a controlled overlay rather than spiked re-calling, and the open-weight arm is a single-replicate pilot whose execution-arm coverage is limited by structured-output and provenance conformance rather than by safety or classification. None of these limits affect the central claim, which is methodological: trustworthiness in agentic genomics is achieved by identifying, measuring and controlling the layers where uncertainty and failure arise.

## Methods

### Overview and design rationale

ClawBench measures *where* uncertainty originates in agentic ACMG/AMP variant interpretation, rather than ranking models by accuracy. A scalar accuracy score conflates distinct failure modes: a variant called “uncertain significance” (VUS) may reflect a safety lapse, missing evidence, unstable code assignment, miscalibrated evidence strength, or a combiner threshold. We therefore assign every variant a per-variant attribution label across an ordered stack of architectural layers, grouped into three blocks: safety control; evidence formation (acquisition, sufficiency, assignment, strength calibration); and decision formation (the deterministic combiner). The framework rests on two methodological pillars described below, temporal blinding of the truth set and a fail-closed evidence contract, which together remove the two confounds that otherwise make agentic interpretation impossible to attribute: pretraining contamination and circularity with the truth label.

### Temporal blinding framework

Large language models may have encountered a variant’s published classification during pretraining, so any benchmark drawn from a public knowledge base must guarantee that the scored label could not have been memorised. We enforce this by temporal blinding: a variant is admissible only if the first-availability date of its current classification strictly post-dates the training cutoff of every model under test, plus a safety margin.

We confirmed the training cutoff of each of the nine candidate models from primary sources (provider documentation and API references), recording a confidence level per model. The binding cutoff was GPT-5.2 at 2025-08-31 (high confidence); the remaining cutoffs ranged from 2024-06-01 to 2025-03-31, all earlier and therefore non-binding. The two lowest-confidence cutoffs sit roughly thirteen months below the boundary and so cannot affect blinding under any plausible correction.

Establishing first-availability is itself a methodological hazard. ClinVar’s date_last_evaluated field records the most recent evaluation, not the first appearance of the current label, and the aggregate variant_summary table ships without submission history [5]. Relying on date_last_evaluated would silently admit variants whose label predates a model’s cutoff. We therefore admit a variant only when first-availability is *provable*: either a dated submission history (reconstructed by joining variant_summary with submission_summary on the variation identifier) places the current tier-group classification after the effective cutoff, or the record’s creation date itself post-dates the cutoff. When neither holds, the variant is excluded (fail-closed).

### Truth-set construction

The truth set, ClinVar release joined to submission-level history [5], was restricted to variants with a review status of at least two stars (multiple submitters, no conflicts, or expert-panel/practice-guideline review). Five-tier classifications were mapped to the canonical ACMG/AMP terms [2]. Streaming over the full variant_summary and submission_summary tables, 166,437 GRCh38 two-star candidates yielded 6,929 temporally blinded variants after the fail-closed first-availability filter removed 159,300 pre-cutoff records and 208 with missing or ambiguous dates (166,437 *−* 159,300 *−* 208 = 6,929); 206 admitted variants had undergone a tier-group reclassification and are flagged. The effective cutoff was 2025-11-29 (binding model cutoff plus a 90-day margin); the earliest admitted label first became available on 2025-11-29, so every label post-dates every confirmed model cutoff by at least 91 days. Temporal blinding controls memorisation of the scored *label*, not of the underlying information: the molecular consequence, population frequency and gene-disease evidence used to classify a variant may predate the cutoff even when the current ClinVar label does not. This is by design, since the benchmark targets reasoning rather than recall, but it means blinding removes one specific channel (label recall) rather than all prior exposure. Sensitivity to the margin is bounded and monotone: the admitted set is 12,464, 10,435, 6,929 and 175 variants at margins of 0, 30, 90 and 180 days respectively; we adopt 90 days. The collapse at 180 days is a window-width artefact, not instability: the ClinVar snapshot was frozen on 2026-06-15 and its admitted labels first became available between 2025-11-30 and 2026-05-27, so a 180-day margin (effective cutoff 2026-02-27) leaves only the roughly three-month window from 2026-02-27 to 2026-05-27, hence 175 variants. Because a margin guards against an *underestimated* binding cutoff, the more consequential error, we also recomputed the set under counterfactual GPT-5.2 cutoffs pushed one, three and six months past the documented 2025-08-31. Blinding is robust to a one- or three-month underestimate: 6,929 and 6,918 of 6,929 labels still post-date the cutoff, and on the Tier-A pilot all 231 variants survive with the dangerous-miscall rate unchanged at 0.7 per cent. Only a six-month underestimate (a true cutoff of 2026-02-28) would erode blinding, leaving the most-recent 175 variants (19 in the pilot); an error of that size would have to exceed the documented cutoff by half a year, and even then admits only the recent-window subset bounded by the snapshot date. The 206 reclassified variants, whose prior label may sit within pretraining, are flagged and stratified separately: of the 14 that fell in the Tier-A pilot, their per-call outcome mix differed from the non-reclassified variants, with more overcalls (40 versus 9 per cent of calls) and a higher dangerous-miscall rate (2.4 versus 0.5 per cent), consistent with genuinely in-flux variants being harder to classify. On 14 variants this is reported as a flag rather than a powered estimate, and it does not affect the layer-attribution results, which are computed on the full panel.

The held-out set was frozen as version 1.0 and is immutable: it carries a content hash over the ordered variant records and a file checksum, and it reproduces bit-for-bit from the released builder. Genomic coordinates are GRCh38 and, where the experiments require them, anchored to Genome in a Bottle high-confidence regions [14]. To ensure the set is not homogeneous, all 6,929 variants were tiered by ACMG automatability from molecular consequence and population allele frequency: Tier A (loss-of-function or common, high expected automatability), Tier B (missense or other protein-altering, moderate), and Tier C (synonymous, intronic or otherwise underdetermined). Tiering localises any performance change to a layer rather than to set composition.

### Attribution framework

#### Constraint gradient

Each variant is interpreted under a constraint gradient that progressively relocates correctness from the stochastic model into executed, validated code: (i) *free* (the model returns a classification from its prior); (ii) *skill-reasoning* (the model reasons over a fixed clinical-variant-reporter specification); and (iii) *skill-execution* (the model returns only structured ACMG evidence codes, which a deterministic engine combines). A truth-supplied condition provides the ceiling control. The truth label never enters any prompt: prompt construction is provably truth-independent, operating only on genomic context and, in the evidence-formation experiments, non-circular evidence.

#### Fail-closed evidence contract

In skill-execution the model’s output is validated against a machine-readable contract before scoring. The contract enforces the ACMG/AMP 28-code vocabulary and strength ladders [2], returns structured errors, and is fail-closed: any violation invalidates the submission rather than defaulting to a permissive interpretation. Critically for blinding, evidence whose provenance is the truth label is blocked. Assertion codes (PP5, BP6) are always disallowed; PS1 and PM5 are disallowed when ClinVar-sourced; and a circularity registry blocks evidence drawn from source families that co-deposit with ClinVar (ClinGen Variant Curation Expert Panels and LOVD). We distinguish ClinGen Sequence Variant Interpretation (SVI) *methods* guidance, which a rigorous submission legitimately cites for strength calibration, from ClinGen *Variant Curation Expert Panel* classifications, which are truth-circular; conflating the two falsely rejects correct in-silico evidence. Strength upgrades above a code’s baseline require an explicit, substantive basis, without which the upgrade is rejected.

#### Deterministic combiners

Validated codes are combined by two deterministic engines used jointly as a decomposition instrument. The first implements the Richards rule-counting logic [2]. The second implements the naturally scaled Tavtigian point system [9]: very-strong and stand-alone evidence score 8 points, strong 4, moderate 2 and supporting 1, with benign codes negated, and the total maps to Pathogenic (*≥* 10), Likely Pathogenic (6 to 9), VUS (0 to 5), Likely Benign (*−*1 to *−*6) and Benign (*≤ −*7). Comparing the two engines on identical codes separates threshold effects (the rule caps where points do not) from evidence effects.

#### Per-variant layer attribution

From the skill-execution replicates of each variant we compute four flags that map to the layers. *safety_clean*: no replicate crosses the lethal Pathogenic-to-Benign boundary relative to truth. *combiner_sensitive*: the modal class from the rule engine differs from the modal class from the point engine on the same codes (a decision-formation effect). *assignment_unstable*: the proposed code sets differ across replicates (an assignment-layer effect). *evidence_insufficient*: the truth is definitive yet even the point engine returns VUS, indicating the model could not form evidence sufficient to classify (a sufficiency or acquisition effect). For the transition analyses we collapse the flags to one ordered category per variant with precedence dangerous *>* evidence_insufficient *>* combiner_sensitive *>* assignment_unstable *>* resolved; VUS-truth controls are categorised separately as correct or overcalled.

### Pilot design

Figure 5 traces how the held-out set feeds each arm described below. The pilot established the baseline decomposition on Tier-A variants. Three temporally compatible models were used: GPT-5.2 (cutoff 2025-08-31), Claude Sonnet 4.5 (2025-03-31) and Gemini 2.5 Flash (2025-01-31), all below the slice boundary. We ran a balanced Tier-A panel of 231 variants (60 Pathogenic, 60 Likely Pathogenic, 60 Benign, 9 Likely Benign and 42 VUS; 14 reclassified) across the free, skill-reasoning and skill-execution conditions, with five replicates per variant and condition. Models were called through a uniform adapter that retries transient and rate-limit errors with exponential backoff and records exhausted rate-limit failures as a distinct category, never as a model format failure, to avoid the silent-throttling artefact that can masquerade as model incompetence.

**Figure 1:**
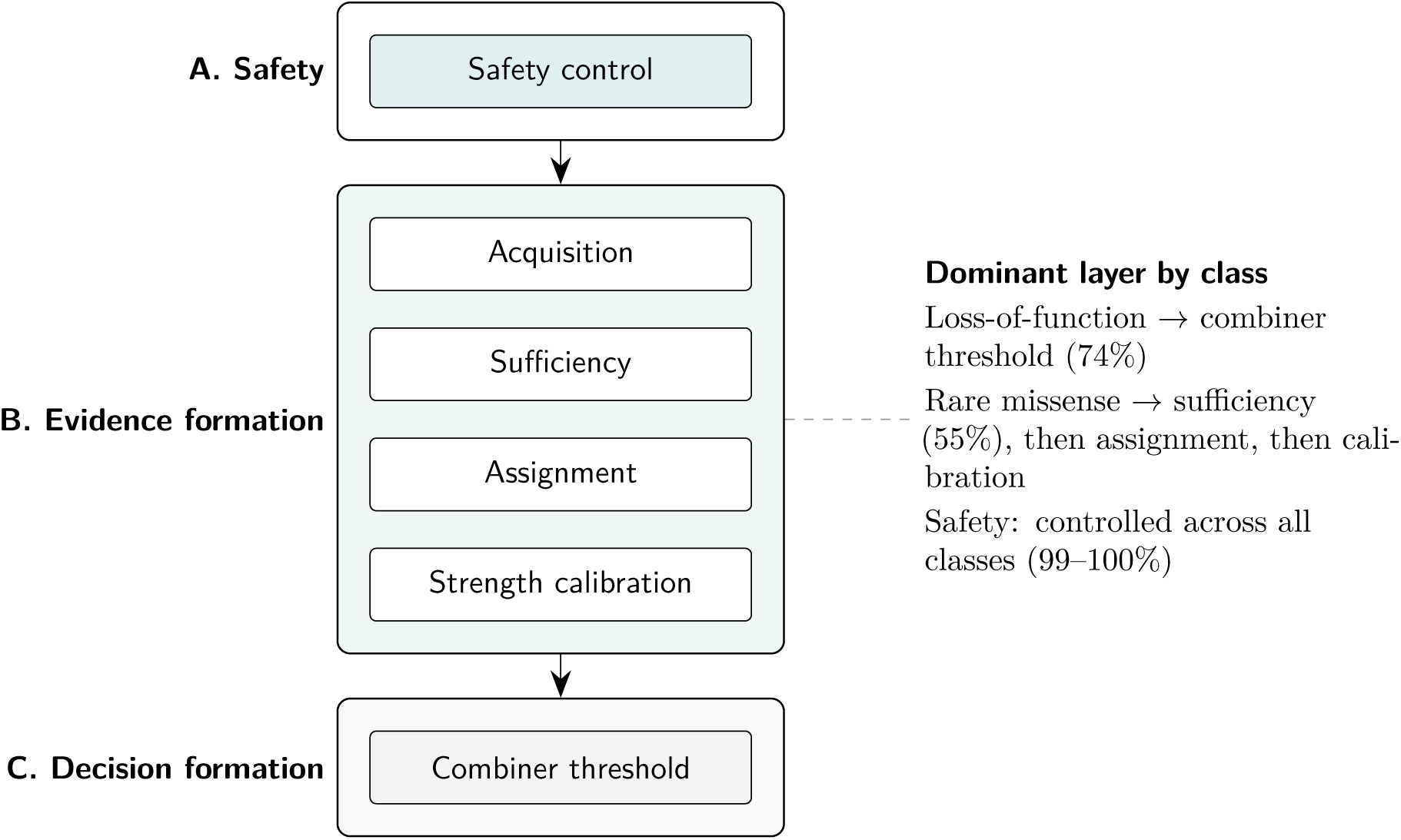
The three-block, six-layer ClawBench interpretation architecture. Safety (block A) is a controlled precondition; evidence formation (block B) carries the four mechanisms where clinical outcomes move; decision formation (block C) is the deterministic combiner. The dominant rate-limiting layer is a property of the variant class.

**Figure 2:**
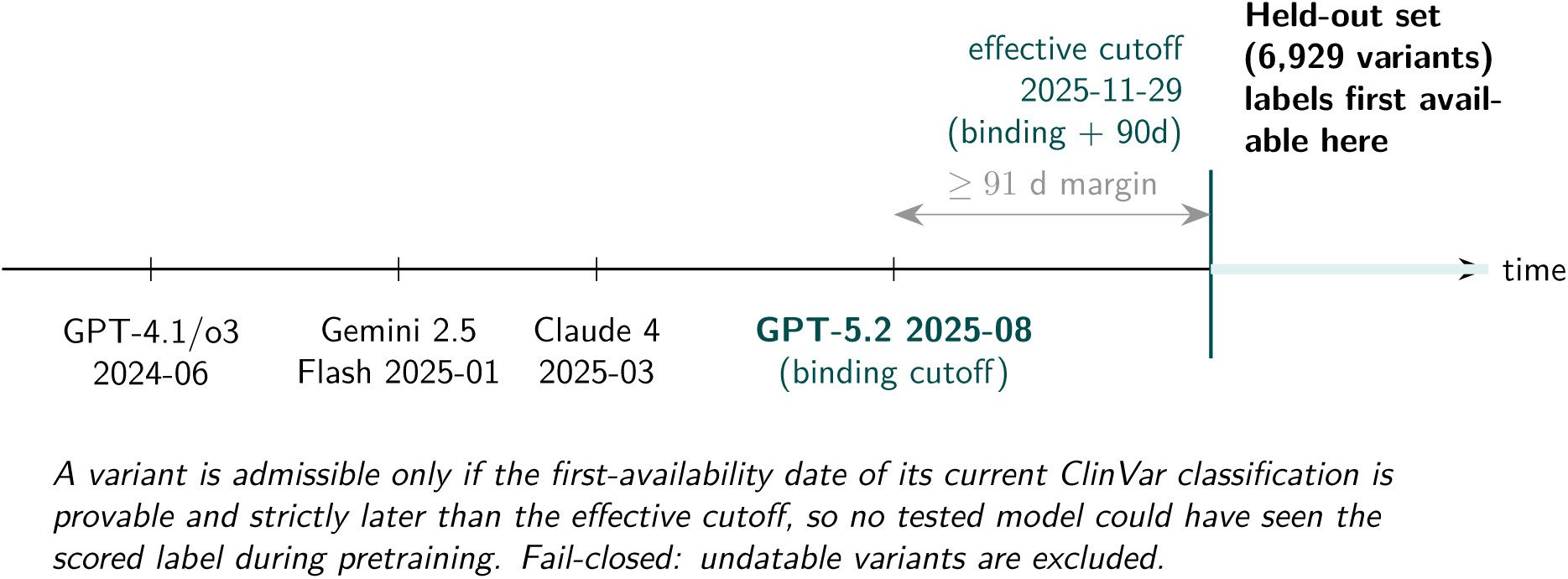
Temporal blinding. Every held-out ClinVar label first became available only after the effective cutoff (the binding model cutoff plus a 90-day margin), so no tested model could have seen the scored label during pretraining. Admission is fail-closed: variants whose first-availability cannot be proven are excluded.

**Figure 3:**
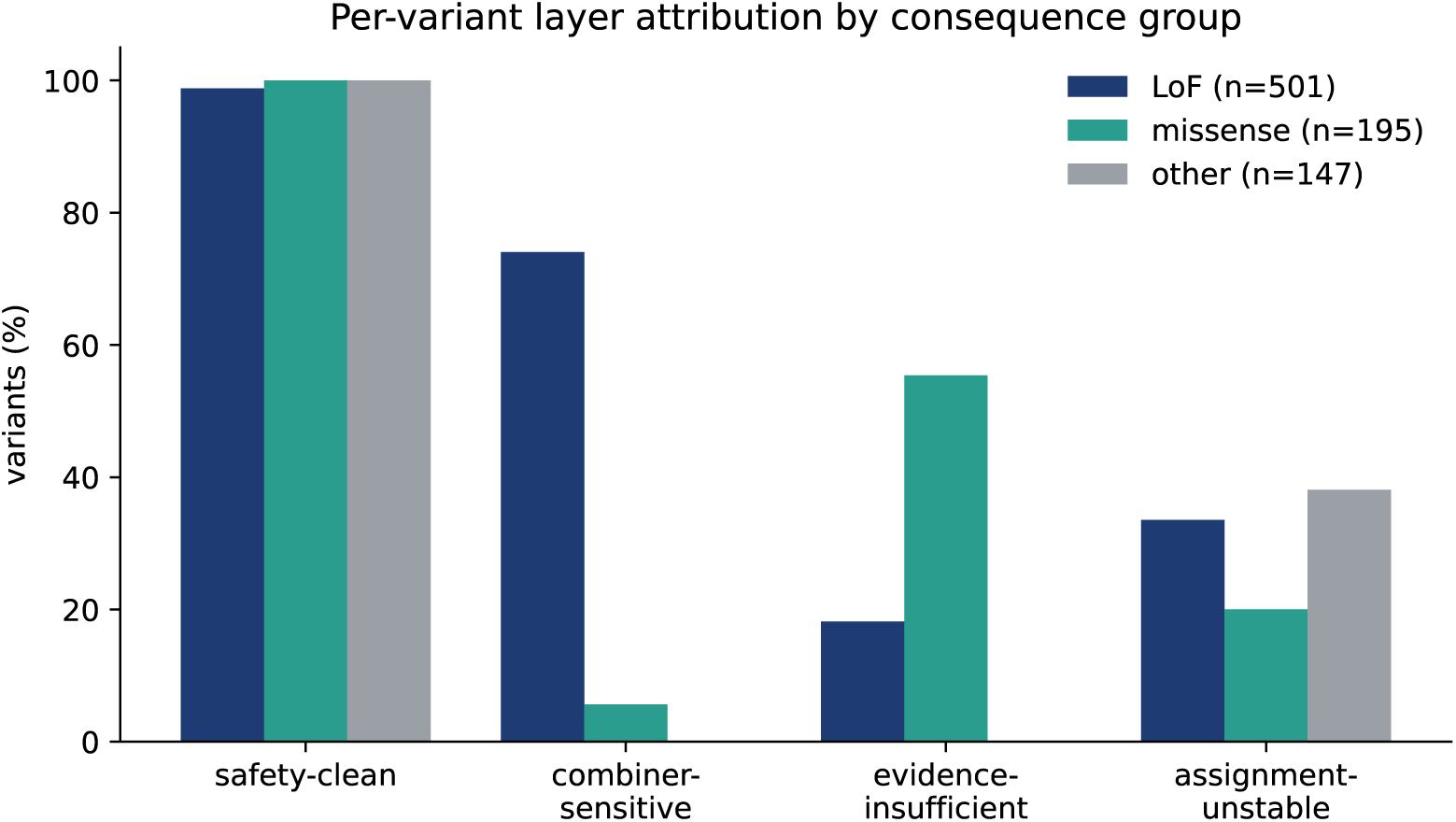
Per-variant layer attribution by consequence group across the skill-execution corpus. Safety-clean is near-universal; combiner sensitivity concentrates in loss-of-function variants, while evidence insufficiency dominates rare missense.

**Figure 4:**
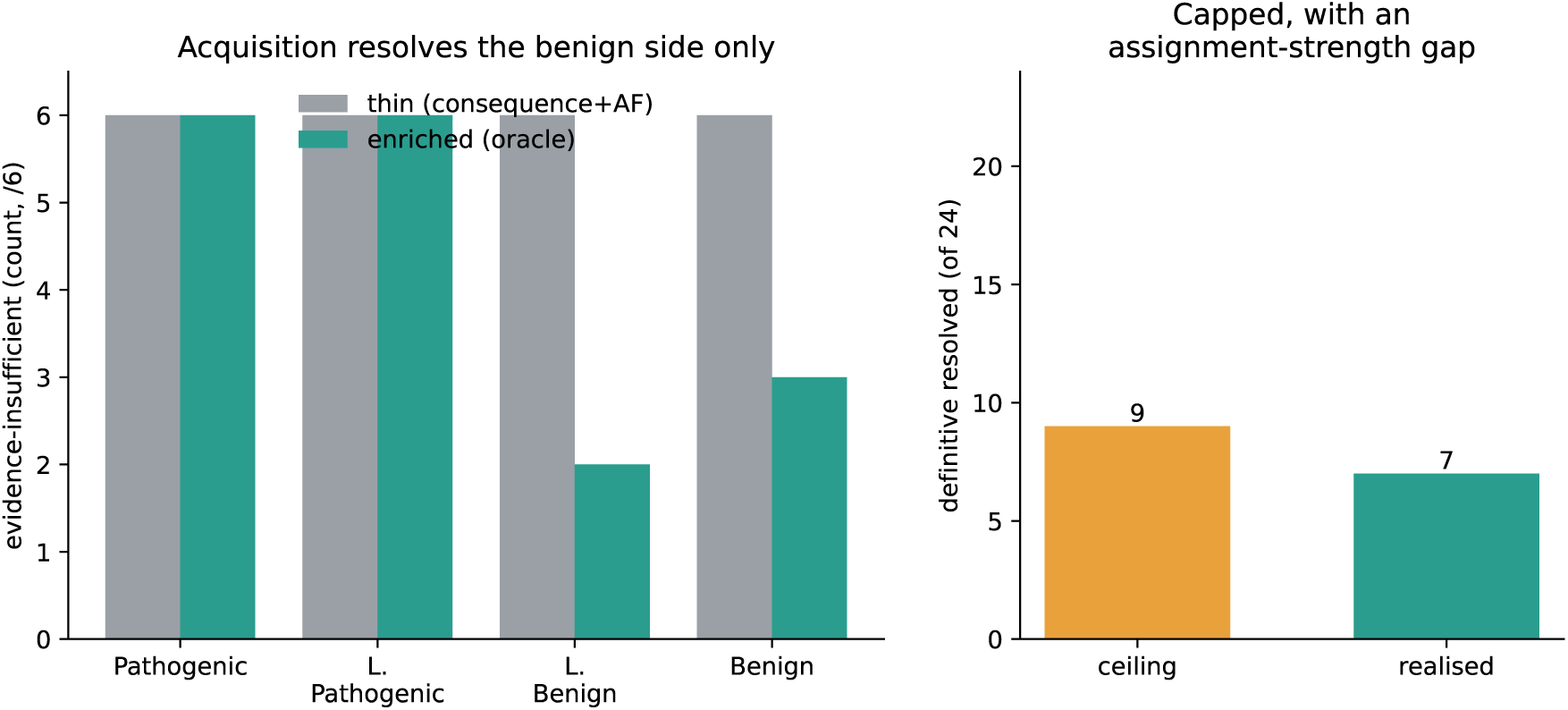
Acquisition arm (Claude Sonnet 4.5, 24 definitive rare-missense variants). Oracle evidence resolves the benign side only, none of the pathogenic side (left); even an ideal agent reaches a ceiling of 9 of 24, and the model realises 7, the gap being an assignment-strength effect (right).

**Figure 5:**
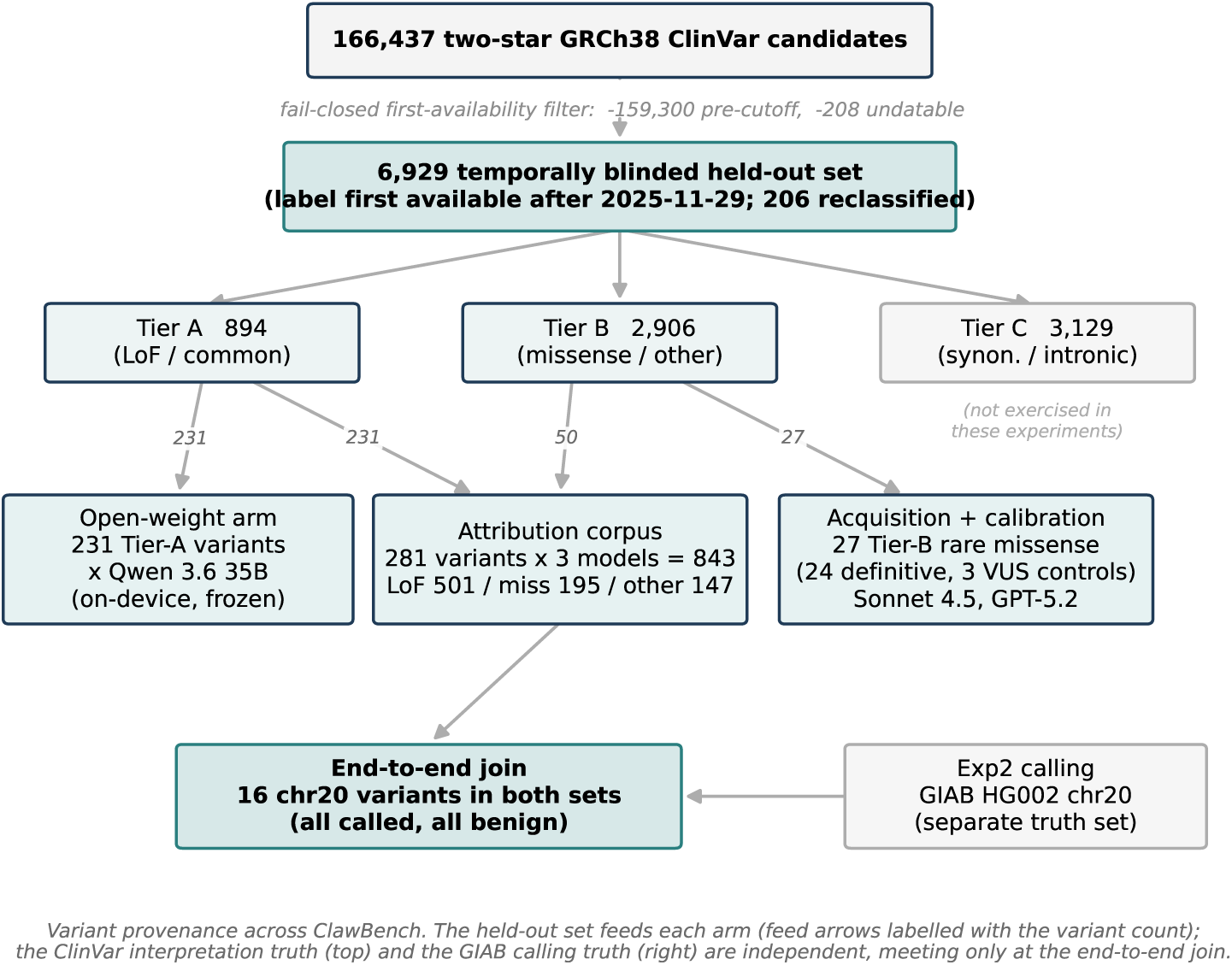
Variant provenance across ClawBench. The temporally blinded held-out set (6,929 variants) is tiered and feeds each experimental arm: the per-variant attribution corpus (281 variants across three models, 843 records; consequence groups loss-of-function 501, missense 195, other 147), the acquisition and calibration arms (27 Tier-B rare missense), and the open-weight arm (231 Tier-A variants). The ClinVar interpretation truth and the independent GIAB calling truth (chromosome 20) meet only at the end-to-end join (16 overlapping variants, all correctly called and all benign). Regenerate with figures/make_flow_figure.py.

### Acquisition-arm design

The acquisition arm tests whether allowing an agent to acquire additional non-circular evidence shrinks the evidence-insufficient layer for rare missense, the class the pilot showed to be dominated by evidence formation. We selected 27 Tier-B rare missense variants by joining the held-out manifest to recover genomic coordinates and gene: 24 definitive variants (six each Pathogenic, Likely Pathogenic, Likely Benign and Benign), enriched for variants flagged evi-dence_insufficient under thin evidence for the model under test, plus three VUS variants as negative controls. The selection is deterministic, frozen and content-hashed.

Each variant was interpreted under skill-execution in two paired conditions differing only in the evidence payload. In the *thin* condition (Condition A, the pilot baseline) the evidence comprised molecular consequence and population maximum allele frequency. In the *enriched* condition (Condition B, an oracle) we deterministically retrieved real non-circular annotations from the Ensembl Variant Effect Predictor REST interface [11], restricted to the canonical or MANE Select transcript: protein consequence and amino-acid change, SIFT, PolyPhen, CADD and AlphaMissense [13], and REVEL [12]. The oracle is the ceiling of acquisition: complete, real evidence, so that a null result cannot be attributed to weak retrieval. ClinVar significance and any colocated-variant clinical-significance fields were scrubbed from the retrieved record before it reached the model, with the fail-closed validator as a second guard against leakage.

The PP3/BP4 evidence line was derived from a single calibrated predictor, REVEL, following ClinGen SVI recommendations not to stack multiple in-silico predictors [15]. We applied the published bidirectional thresholds: PP3 at supporting, moderate and strong for REVEL at least 0.644, 0.773 and 0.932 respectively, and BP4 at strong for REVEL at most 0.016 and at supporting for REVEL at most 0.290; the published benign-moderate band was collapsed into BP4 supporting because the 2015 framework has no benign-moderate level, and scores between 0.291 and 0.643 license no PP3/BP4. The other predictors were supplied as corroborating context only. Because applying a calibrated strength above the supporting baseline requires an explicit basis under the contract, the enriched payload provided the calibration citation as the strength basis. The model under test was Claude Sonnet 4.5, with five replicates per variant and condition.

### Calibration-arm design

The acquisition arm localised the residual gap to strength assignment: the model applied PM2 at supporting rather than at its 2015 moderate baseline, leaving rare pathogenic missense one point below Likely Pathogenic even with a strong calibrated PP3. The calibration arm tests whether explicit strength-calibration guidance recovers the ceiling. It re-ran the identical 27 variants and oracle evidence under skill-execution, adding a single instruction to the enriched payload: that this benchmark scores by ACMG 2015 strength baselines, so PM2 applies at moderate when warranted, rather than the ClinGen SVI 2020 downgrade to supporting. The instruction states the scoring convention; it does not dictate code direction or whether PM2 applies. Five replicates per variant were run for Claude Sonnet 4.5, and the same three-arm protocol (thin, enriched, calibrated) was repeated for GPT-5.2 as a confirmatory model to test whether any calibration effect is model-general.

### Statistical analysis

Primary endpoints are per-variant attribution categories, computed deterministically from the five skill-execution replicates of each variant and arm as described above. We report, for each arm, the count of definitive variants in each attribution category, and the thin-to-enriched and enriched-to-calibrated category transition matrices. The headline acquisition endpoint is the proportion of definitive variants that leave the evidence_insufficient category between arms.

To separate what is achievable from what was achieved, we compute a deterministic theoretical ceiling per variant: the best point-engine class reachable from the oracle’s non-circular evidence under an ideal, direction-aware strength assignment (PM2 at its moderate baseline plus the calibrated PP3 on the pathogenic side, or the calibrated BP4 alone on the benign side, with no contradictory PM2). The gap between the ceiling and the realised count quantifies assignment-strength conservatism.

Replicate agreement is reported per cell; submissions failing the fail-closed contract are recorded as invalid with their error codes and excluded from attribution, and the invalid rate is reported per arm. Because the evidence-formation arms involve 24 definitive variants and the calibration effect arises from deterministic combiner arithmetic rather than from sampling variation, we report counts and proportions rather than inferential test statistics for these arms, and establish robustness through two-model confirmation rather than through a single-model significance claim; the pilot endpoints, computed over the larger Tier-A panel, are reported with case-level uncertainty. All quantitative claims are regenerated from released per-variant data by the analysis scripts, and the held-out truth set, frozen probe set and oracle evidence cache are content-hashed for exact reproducibility.

### Execution benchmark: the calling constraint gradient

The execution half evaluates germline short-variant calling from paired sequencing reads against Genome in a Bottle (GIAB) truth sets [14]. Four arms process the same sample: (A) a free agent given the task and read paths only; (B) a skill-reasoning agent additionally given the wrapper specification; (C) a skill-execution agent that invokes the validated, version- and container-pinned nf-core/sarek wrapper [16; 17] running GATK HaplotypeCaller [18; 19]; and (D) a hand-run best-practice reference. Arms A and B execute the workflow the agent authored, inside a sandbox with read-only inputs, no network writes and resource limits, so each run fails or succeeds on its own merits. Calls are scored against the GIAB confident regions with rtg vcfeval [20] following GA4GH benchmarking best practice [21].

### Pre-registered failure taxonomy and trust-property scorecard

Before any agent arm was scored we pre-registered eight failure labels and a deterministic classifier that assigns them from run artefacts (the emitted command, the provenance manifest, the produced VCF, a two-run genotype comparison and vcfeval output): reference-build error, container-version error, parameter drift, missing provenance, non-reproducibility, accuracy degradation, incomplete workflow and tool-selection error. A run is clean only if it carries none. The per-run trustworthiness scorecard records five dimensions, accuracy (vcfeval F1 within the confident region), reproducibility (genotype-identical re-run), provenance (a complete manifest), pinning (digest-pinned containers and pinned versions) and taxonomy cleanliness; a run is trustworthy only if it satisfies all five. Plan validity is scored separately at the command level: the emitted workflow must contain the mandatory stages (alignment and calling, or an encapsulated validated pipeline) in canonical order, judged from the command text and not from convergence.

### Open-weight arm

To make the benchmark reproducible without a paid endpoint, the interpretation gradient was additionally run on a local open-weight model (Qwen, served on-device through a local inference server) on the identical temporally blinded variants and contract used for the frontier models. Open weights are frozen and hash-identifiable, so the open-weight arm reproduces exactly; the model adapter is the only addition, the scoring path is unchanged.

### End-to-end attribution

For variants that are both GIAB-confident truth calls and members of the held-out interpretation set, the per-variant calling outcome (true positive with matching genotype, true positive with a wrong genotype, or false negative) is joined to the interpretation attribution to yield a single end-to-end label. False negatives are attributed to the calling layer and never reach interpretation; a wrong genotype is a calling error that propagates into interpretation; a correctly called variant is attributed to its rate-limiting interpretation layer, or flagged as a dangerous misclassification. Because reference genomes are benign-dominated, pathogenic end-to-end cases are exercised with a controlled overlay that conditions on correct calling (the validated arm calls GIAB truth at F1 0.99) and applies the real interpretation attribution; this is a labelled positive control demonstrating the joint attribution, not a measurement of caller performance on spiked variants.

### Scope

This study is scoped to the measurement framework. Calling accuracy is reported as qualification of the validated control arm on a chromosome-20 development pilot, not as a genome-wide benchmark result; cross-host reproducibility beyond the same-host genotype-identical re-run, and multi-replicate variance for the open-weight arm, are stated as limitations rather than claimed.

## Data and code availability

The ClawBench harness, the deterministic classifiers and the analysis scripts are released at https://github.com/ClawBio/ClawBench (MIT licence), the public repository that accompanies this preprint. The temporally blinded held-out truth set is immutable and content-hashed; the manifest carries the SHA-256 content hash 8d745d32ad8a377f05022174f80041d0f237aa24fc5c1dcb87fcebe16c242c66. The held-out manifest, the agent-arm emissions and the per-ancestry calling table are being deposited as a versioned reproduction archive (Zenodo). We assert no reproduction beyond these released artefacts and their content hashes; a clean-clone rebuild and the archival deposit will accompany the journal submission.

